# Visual imagery and visual perception induce similar changes in occipital slow waves of sleep

**DOI:** 10.1101/532317

**Authors:** Giulio Bernardi, Monica Betta, Jacinthe Cataldi, Andrea Leo, José Haba-Rubio, Raphael Heinzer, Chiara Cirelli, Giulio Tononi, Pietro Pietrini, Emiliano Ricciardi, Francesca Siclari

## Abstract

Previous studies have shown that regional slow wave activity (SWA) during NREM-sleep is modulated by prior experience and learning. While this effect has been convincingly demonstrated for the sensorimotor domain, attempts to extend these findings to the visual system have provided mixed results. Here we asked whether depriving subjects of external visual stimuli during daytime would lead to regional changes in slow waves during sleep and whether the degree of ‘internal visual stimulation’ (spontaneous imagery) would influence such changes. In two 8h-long sessions spaced one-week apart, twelve healthy volunteers either were blindfolded while listening to audiobooks or watched movies (control condition), after which their sleep was recorded with high-density EEG. We found that during NREM-sleep the number of small, local slow waves in the occipital cortex decreased after listening with blindfolding relative to movie watching in a way that depended on the degree of visual imagery subjects reported during blindfolding: subjects with low visual imagery showed a significant reduction of occipital sleep slow waves, while those who reported a high degree of visual imagery did not. We also found a positive relationship between the reliance on visual imagery during blindfolding and audiobook listening and the degree of correlation in sleep SWA between visual areas and language-related areas. These preliminary results demonstrate that short-term alterations in visual experience may trigger slow wave changes in cortical visual areas. Furthermore, they suggest that plasticity-related EEG changes during sleep may reflect externally induced (‘bottom-up’) visual experiences, as well as internally generated (‘top-down’) processes.

## Introduction

Slow waves represent the main electroencephalographic (EEG) signature of non-REM (NREM)-sleep in humans, as well as in most mammals and birds (Cirelli & Tononi, 2008). They occur when neurons become *‘bistable’* and undergo a slow oscillation in membrane potential between a hyperpolarized ‘silent’ phase (*down-state*) and a depolarized period characterized by a high firing activity (*up-state*) (Steriade et al., 1993). Importantly, the EEG signal power in the 0.5–4.0 Hz (*delta*) range, referred to as *‘slow wave activity’* (SWA), has been shown to represent a reliable marker of homeostatically regulated sleep need. In fact, SWA shows its maximum during the first hours of sleep, decreases progressively in the course of the night, and increases proportionally with the duration of prior wakefulness and after periods of sleep restriction (Achermann & Borbély, 2003). Recent work has demonstrated that changes in SWA are not uniformly distributed across the cortex. Instead, slow waves occur and are regulated locally (Nir et al., 2011), in an experience-dependent manner (Siclari & Tononi, 2017). For example, brain areas that are intensely recruited during a visuo-motor task typically show increased SWA during subsequent sleep (Huber et al., 2004). Conversely, prolonged arm immobilization results in local decreases in SWA over sensorimotor cortex (Huber et al., 2006). In addition, local changes in SWA within sensorimotor areas correlate with post-sleep improvement in visuo-motor performance (Fattinger et al., 2017). Animal studies and computer simulations suggest that local SWA changes may both reflect experience-dependent changes in regional synaptic density and strength and play a role in cellular and systems restoration through synaptic renormalization (Tononi & Cirelli, 2014). However, changes in SWA may also reflect changes in on the neuromodulatory tone and the balance between excitation and inhibition, independent of changes in synaptic strength (Cirelli, 2017; Frank & Cantera, 2014). Moreover, the wake-dependent accumulation of metabolites and of molecular factors reflecting an increased cellular stress have been also suggested to potentially affect SWA at both global and local level (Krueger et al., 2008; Qi et al., 2016; Vyazovskiy & Harris, 2013; Wang et al., 2018).

It is currently less clear whether the visual system is similar to the sensorimotor system in the way sleep SWA is modulated based on waking experiences. In particular, two recent studies employing a visual perceptual learning paradigm (Bang et al., 2014; Mascetti et al., 2013), did not find significant changes in occipital SWA during the post-training night, although one of them (Mascetti et al., 2013) revealed a positive correlation between the number of slow waves initiated in a task-related occipito-parietal area and post-sleep performance improvement. Another study (Korf et al., 2017) reported a relative reduction in SWA following visual deprivation, but failed to show a regionally specific effect in visual cortical areas. It should be noted, however, that these studies categorized slow waves based on parameters that may not be fully appropriate for occipital (visual) areas. For example, compared to other cortical regions, occipital slow waves are typically smaller and show smaller modifications during recovery sleep after sleep deprivation/restriction (Finelli et al., 2001). In addition, recent work has provided evidence for two types of slow waves (widespread, *‘type I’* and local, *‘type II’*) that are synchronized through different mechanisms and have distinct spatiotemporal characteristics, and which should therefore be analyzed separately (Bernardi et al., 2018; Siclari et al., 2014). Finally, changes in slow wave parameters after visual deprivation could be affected by the occurrence or not of mental imagery, which involves the generation of endogenous, ‘quasi-perceptual’ sensory experiences through a ‘*top-down*’ activation of visual cortical areas (Dentico et al., 2014). Indeed, there is evidence that imagery-related experiences may lead to plastic adaptations similar to those induced by actual perceptual stimuli or real actions (e.g., Jackson et al., 2003; Pascual-Leone et al., 1995).

In this preliminary study, we set out to investigate local effects of short-term visual deprivation on slow waves during subsequent NREM-sleep using high-density (hd-)EEG recordings, which combines the high temporal resolution of standard EEG and an improved spatial resolution through source reconstruction techniques. Moreover, we assessed potential changes of both small/local slow waves and large/widespread slow waves. Finally, we investigated whether ‘internal visual stimulation’ (mental imagery) would influence slow wave changes induced by visual deprivation.

## Methods

### Subjects

Twelve healthy volunteers (age 25.5 ± 3.7 yrs, 6 F) screened for neurological, psychiatric, and sleep disorders and who were not on psychotropic medication participated in the study. The sample size has been determined based on previous investigations regarding local, experience dependent changes in slow wave characteristics within the sensorimotor domain (Huber et al., 2006, 2004; Kattler et al., 1994). All these studies employed an experimental paradigm similar to the one used in present work and included a similar number of subjects (range 8-14). A power calculation performed using the smallest effect size reported across these investigations (0.94; Kattler et al., 1994) and a desired statistical power of 0.8, indicated a sample size of 11 as sufficient to detect a similar or stronger effect.

All volunteers had a good sleep quality as assessed by the *Pittsburgh Sleep Quality Index* (PSQI score ≤ 5; Buysse et al., 1989), and scored less than 10 points on the *Epworth Sleepiness Scale* (ESS; Johns, 1991). None of the participants had an extreme chronotype, as determined using the *Morningness-Eveningness Questionnaire* (Horne & Ostberg, 1975). Moreover, all participants showed intermediate levels of visual imagery skills (e.g., Zeman et al., 2015), with scores in the *Vividness of Visual Imagery Questionnaire* (VVIQ; Marks, 1973) comprised between 49 and 69, mean 57.2 ± 6.4.

Volunteers were asked to maintain a regular sleep–wake schedule in the five days preceding each experiment. Compliance was verified by wrist-worn actigraphy devices (MotionWatch 8, CamNtech; also see Table 1 for quantitative comparisons). The study was approved by the ethical committee of the Lausanne University Hospital. Written informed consent was obtained from each subject.

**Table 1.**
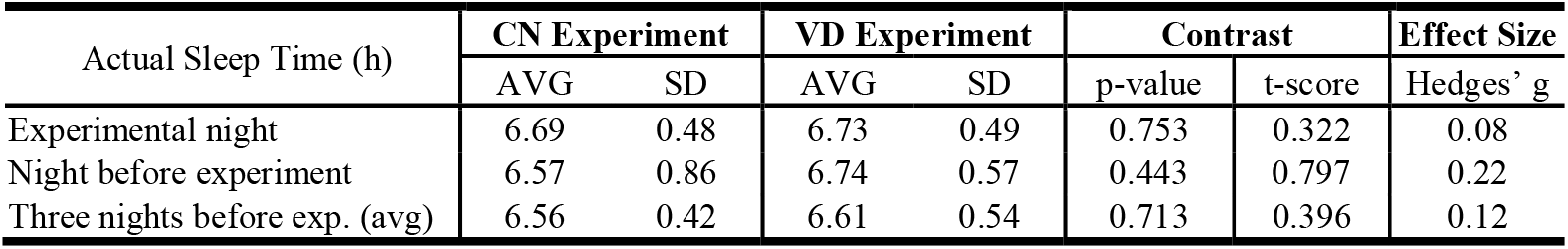
Actigraphy results. Mean ‘actual sleep time’ measured before each experiment and during the experimental night (AVG, average; SD, standard deviation). Values obtained during the experimental night were similar to those recorded at home, either the night before the experiment or in the three days (average) before the visual deprivation (VD) and control (CN) experiments. No significant effects were observed (CN vs. VD contrast and experimental-night vs. nights-at-home contrast; paired t-tests; p > 0.05).

| Actual Sleep Time (h) | CN Experiment |  | VD Experiment |  | Contrast |  | Effect Size |
| --- | --- | --- | --- | --- | --- | --- | --- |
|  | AVG | SD | AVG | SD | p-value | t-score | Hedges' g |
| Experimental night | 6.69 | 0.48 | 6.73 | 0.49 | 0.753 | 0.322 | 0.08 |
| Night before experiment | 6.57 | 0.86 | 6.74 | 0.57 | 0.443 | 0.797 | 0.22 |
| Three nights before exp. (avg) | 6.56 | 0.42 | 6.61 | 0.54 | 0.713 | 0.396 | 0.12 |

### Experimental Design

Participants completed two experimental sessions, separated by at least one week and performed in a random and counterbalanced order: a visual deprivation (VD) condition, during which subjects were blindfolded, and a control (CN) condition. In both conditions, participants arrived at the sleep laboratory at ~2 PM, were assigned to a dedicated experimental room, and were given ~45 min to familiarize themselves with the environment. In the VD condition, the room was kept dark and subjects were blindfolded with a sleeping masks and opaque eye patches for the whole duration of the experiment. All wake activities during the two experiments were rigorously controlled: during VD, subjects had to listen to audiobooks during three ~2 h periods (~6 h in total), while during CN they watched movies (with audio) for a similar amount of time (Figure 1). To enhance compliance, subjects were allowed to select the material for both conditions from a pre-defined list including adventure and fantasy stories without strong emotional contents. Between 9 PM and 10.30 PM, the hd-EEG sensor-net (256 channels, 500 Hz sampling frequency; Electrical Geodesics Inc.) was applied and the spatial coordinates of all electrodes were registered using the Geodesic Photogrammetry System (GPS 3.0). To ensure optimal signal quality during EEG recordings, electrode impedance was checked before each recording and kept below 50 KΩ. All participants were allowed to sleep for ~7.5 h (11.30 PM – 7.00 AM). An adaptation night was not performed.

**Figure 1.**
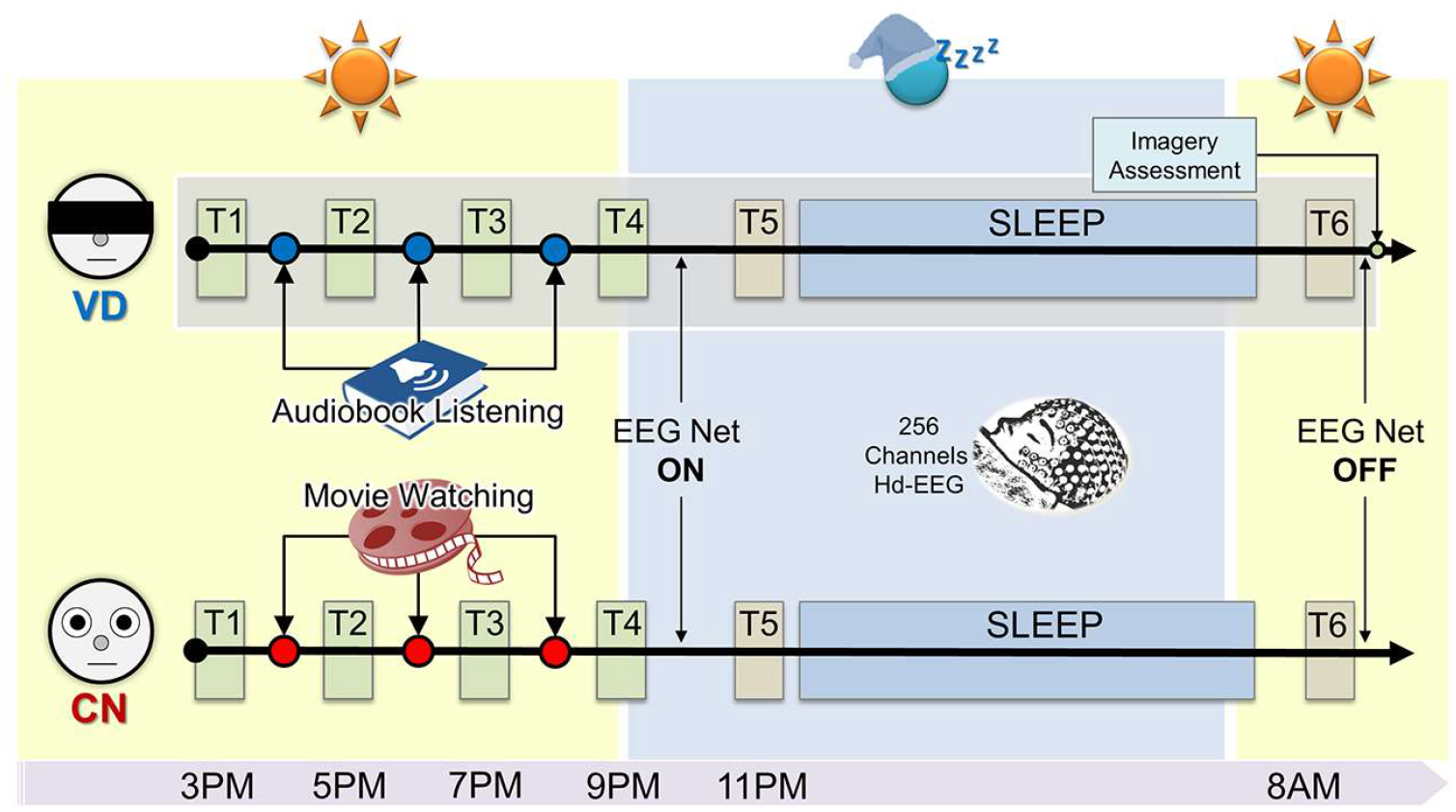
Experimental paradigm. Each test block (T1-T6) included an auditory psychomotor vigilance test (aPVT) and Likert scales for sleepiness, alertness and mood. Sessions T5 (before sleep) and T6 (after sleep) also included three 2 min EEG recording sessions with eyes closed. During the visual deprivation (VD) experiment participants were blindfolded and listened to audiobooks for a total of ~6 h (three 2 h sessions), while in the control (CN) experiment they watched movies for a similar amount of time. Brain activity during sleep was recorded using hd-EEG.

Brief test sessions including an auditory psychomotor vigilance test (aPVT, see below; Jung et al., 2011) and Likert-scales (range 1-10) for sleepiness, alertness and mood were completed every 2 h (T1-T5), and in the morning, after awakening (T6). The morning test session was initiated at least 30 min after awakening (~8 AM) to minimize the possible influence of sleep inertia (Jewett et al., 1999). Three 2-min (6 min in total) eyes closed hd-EEG recordings were obtained before (T5) and after sleep (T6), while the subjects lied still and awake in bed. At the end of the VD experiment, participants completed a short debriefing questionnaire including Likert-scales (range 1-5) aimed at evaluating how much they had relied on visual imagery during non-visual (i, auditory; ii, tactile) experiences (*“How much did you rely on visual imagery during auditory/tactile experiences?”*). In order to estimate the general tendency of each subject to rely on visual imagery during non-visual experiences, an ‘overall’ imagery score was calculated by averaging scores reported for auditory and tactile experiences. The possible occurrence of hallucinatory episodes was also investigated.

Standardized quantities of food were provided at ~6:00 PM and at ~7:15 AM (after sleep) the following day. Alcohol-containing beverages were prohibited starting the night before the experiment and throughout each experiment. Subjects were asked to restrain from assuming caffeine-containing beverages during the two experiments. In order to avoid excessively long periods of immobility, volunteers were allowed to take short breaks between audiobook-listening and movie-watching sessions. Experimenters took turns at monitoring the participants to prevent them from falling asleep and to ensure adherence to the protocol throughout each experiment. Moreover, after each session of audiobook listening or movie watching, a brief oral summary was requested from all volunteers to verify attention to the task.

### Auditory PVT and Subjective Scales

During the aPVT (adapted from Hung et al., 2013, generated using E-Prime 2, Psychology Software Tools, Inc.), volunteers were instructed to respond as fast as possible (by pressing a button) to a continuous 1000 Hz tone. Tones were presented binaurally at an unchanging, comfortable volume (~45 dB) through earbuds while subjects sat still with their eyes closed. Following each button press (or after 5 s if no response was produced) the tone was stopped and a new sound was presented after a randomly selected interval comprised between 2 and 12 s. The total duration of each test trial was ~10 min. The mean reaction time during the aPVT was used as an objective measure of the participants’ vigilance level (reaction times > 500 ms were considered as lapses).

Specific analyses were used to verify whether experimental procedures in CN and VD had comparable effects on aPVT reaction time and subjective measures of vigilance, sleepiness and mood. Specifically, changes in these scores at 11 PM (before sleep, after ~8 h of visual deprivation or stimulation) were calculated with respect to the first measurement, performed at 3 PM. This variation was compared across the two experimental conditions (CN, VD) using paired t-tests.

### Wake EEG Recordings

Previous work showed that prolonged practice with particular tasks may lead to a local, regionally-specific increase in low-frequency activity during wakefulness in the theta range (5-9 Hz) (Hung et al., 2013), which may reflect experience-dependent changes in sleep need similar to local SWA during NREM-sleep (Bernardi et al., 2015, 2016; Hung et al., 2013; Nir et al., 2017; Pigarev et al., 1997). Thus, we also investigated regional (occipital) changes in *theta* activity during wakefulness after short-term visual deprivation.

For each recording session (before/after sleep), 6 min of spontaneous eyes-closed EEG data were band-pass filtered between 0.5 and 45 Hz. Each recording was divided into non-overlapping 5 s epochs and visually inspected to identify and reject bad channels and epochs containing clear artifacts (NetStation 5, Electrical Geodesic). Then, an Independent Component Analysis (ICA) was performed to remove residual ocular, muscular, and electrocardiograph artifacts using EEGLAB (Delorme & Makeig, 2004). This procedure allows to significantly reduce the impact of EEG artifacts on power computation and on the identification of individual graphoelements, while producing negligible changes in physiological signals of interest (Iriarte et al., 2003; Romero et al., 2003). Rejected channels were interpolated using spherical splines. Finally, the signal of each electrode was re-referenced to average reference. For each EEG derivation, power spectral density (PSD) estimates were computed with the Welch’s method in artifact-free 5 s data segments (Hamming windows, 8 sections, 50% overlap), and integrated between 5 and 9 Hz (*theta* activity).

### Sleep EEG Recordings

Sleep EEG recordings were scored according to standard criteria in 30 s epochs (Iber, 2007). For scoring purposes, electrodes located in the chin-cheek region were used to evaluate muscular activity, while four electrodes placed at the outer canthi of the eyes were used to monitor eye movements (Siclari et al., 2014). Recordings were band-pass filtered between 0.5 and 45 Hz. All epochs scored as N2 or N3 (NREM) were then extracted and visually inspected to remove bad channels (later replaced by spherical spline interpolation). An ICA-based procedure was used to remove potential artifacts, as described above.

Previous work showed that experience-dependent changes in SWA are strongest during the first 20-30 min of NREM-sleep, and tend to dissipate later (Huber et al., 2006, 2004). Thus, here we specifically focused our analyses on the first 20 min of stable (N2/N3) NREM-sleep (EP1). The second 20 min epoch (20-40 min of NREM-sleep; EP2) was also extracted for comparison. Periods containing clear microarousals (duration 3-15 s) or wake periods (as defined during the scoring procedure) were excluded from analysis.

For each electrode, the signal was re-referenced to average reference and SWA activity was calculated as the signal power in the 0.5-4.0 Hz range (using the same procedure described for *theta* activity). Moreover, an automated detection algorithm based on zero-crossings of the signal (Riedner et al., 2007) was used to identify individual slow waves in each electrode. The signal of each channel was referenced to the average of the two mastoid electrodes, down-sampled to 128 Hz and band-pass filtered (0.5-4.0 Hz, stop-band at 0.1 and 10 Hz) before application of the algorithm. Only slow waves with a duration of 0.25-1.0 s between consecutive zero crossings (half-wave) were further examined for the estimation of slow wave density (number of detections per minute; sw/min) and mean negative amplitude (μV). Of note, occipital EEG slow waves are significantly smaller than slow waves originating in other brain regions (Finelli et al., 2001), and large slow waves volume-conducted (or travelling; Massimini et al., 2004) from other areas could thus mask region-specific, occipital changes. Therefore, an additional analysis was performed to further investigate the potential effects of visual deprivation on the density of large and small slow waves. Specifically, for each subject and electrode, an amplitude threshold was applied. The threshold was defined as the mean plus 2 times the mean absolute deviation (MAD) of slow wave amplitude values obtained from the first 20 min of NREM-sleep in the CN night (used as a reference condition): slow waves with an amplitude lower than this threshold were classified as *‘small’*, while the remaining waves were classified as *‘large’*.

### ROI-Based Analysis

Given the specific hypotheses of the present study, analyses were initially focused on two symmetrical regions of interest (ROIs), each including 18 electrodes (Figure 3A): *i*) an occipital (*test*) ROI, centered on Oz and extending to occipito-temporal and occipito-parietal electrodes, and *ii*) a centro-frontal (*control*) ROI, centered on Fz. This control ROI was selected for two main reasons: *i*) it is relatively distant from the occipital area, and thus can be expected to present minimal cross-regional signal contamination caused by volume-conduction; *ii*) it corresponds to one of the scalp areas that is strongly influenced by ‘global’ homeostatic changes in sleep pressure (Finelli et al., 2000; Leemburg et al., 2010), and its evaluation may thus allow to better distinguish between local and global changes in slow wave parameters. Of note, since the sole spatial distance can be expected to not completely eliminate cross-regional EEG-signals contamination due to volume conduction (Jackson & Bolger, 2014), effects in the two ROIs were not directly compared. Instead, a source modeling analysis was planned to verify the regional specificity of potential findings.

### Source Modeling Analysis

In order to better evaluate the spatial distribution of potential changes in slow wave characteristics, a source localization analysis of the 0.5-4.0 Hz band-pass filtered signal was performed on the first 20 min of NREM data using GeoSource 3.0 (NetStation, Electrical Geodesics, Inc). A four-shell head model based on the Montreal Neurological Institute (MNI) atlas and a subject-specific coregistered set of electrode positions were used to construct the forward model. The inverse matrix was computed using the standardized low-resolution brain electromagnetic tomography (sLORETA) constraint, since previous studies showed that this approach is well-suited to identify local aspects of sleep slow waves at the group-level (Anderer & Saletu, 2013; Castelnovo et al., 2016; Mander et al., 2015; Mander et al., 2011; Murphy et al., 2009; Riedner et al., 2011; Saletin et al., 2013; Siclari et al., 2014). The source space was restricted to 2,447 dipoles distributed over 7 mm^3^ cortical voxels and a Tikhonov regularization procedure (λ = 10^−2^) was applied to account for the variability in the signal-to-noise ratio. Since a slow wave detection procedure cannot be applied to this signal, the average standard deviation (SD) of current source density time-series was used as a proxy for mean slow wave amplitude at each voxel. Indeed, the standard deviation of the 0.5-4.0 Hz signal reflects the extent to which the peaks and troughs of signal oscillations differ from the mean current density, and thus provides indirect information about the overall amplitude of slow waves.

### Statistical Analysis

Direct statistical comparisons between the two experimental conditions (CN, VD) were performed using two-tailed paired parametric t-tests (N=12 per condition). Comparisons involving only a subsample of study participants (N=6 per group/condition) were computed using non-parametric tests for paired (Wilcoxon signed-rank test) or unpaired (Mann–Whitney U test) samples. The role of visual imagery was investigated using Spearman’s correlation (N=12). A bias-corrected and accelerated bootstrapping procedure (BCa 95%, 5000 iterations; Davison & Hinkley, 1997) was used to estimate the confidence intervals of paired comparisons and correlations performed at ROI-level. One-tailed tests were used for analyses based on strong a-priori assumptions regarding the specific direction (positive or negative) of the comparison/correlation (Figure 7A and Figure 8). For analyses testing the same hypothesis on multiple electrodes/voxels, a correction for multiple comparisons was performed using a False Discovery Rate (FDR) adjustment (Benjamini & Hochberg, 1995).

## Results

### Effects of visual deprivation or stimulation on vigilance, mood and sleep architecture

Participants completed two experimental sessions spaced at least one week apart: in VD (visual deprivation) subjects were blindfolded and listened to audiobooks, and in CN (control condition) they watched movies. At the end of the 8h waking period of VD or CN, before sleep (11 PM), subjects showed similar relative variations in aPVT reaction time (used as an index of vigilance level; p = 0.195, t_(11)_ = 1.379), and in subjective sleepiness (p = 0.359, t_(11)_ = 0.957), alertness (p = 0.096, t_(11)_ = 1.821) and mood (p = 0.085, t_(11)_ = 1.893; Figure 2).

**Figure 2.**
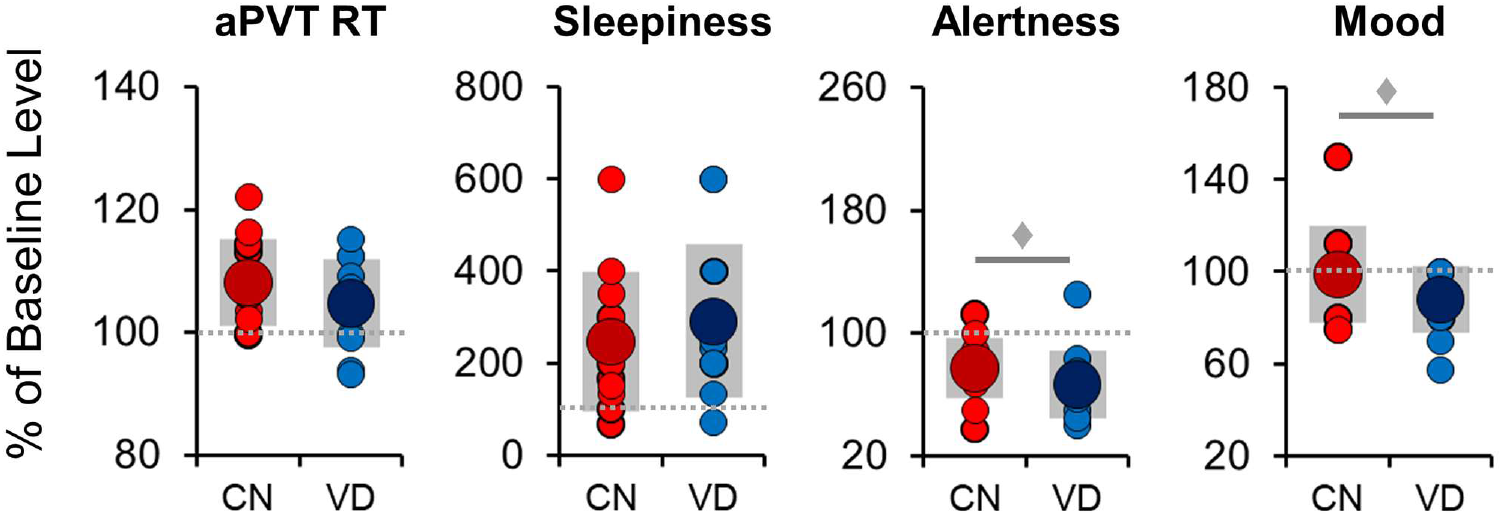
Changes in vigilance and mood during experiments. Relative changes in auditory psychomotor vigilance test (aPVT) reaction time, sleepiness, alertness and mood induced by experimental procedures (i.e., after 8 h of visual stimulation or visual deprivation; 11 PM). Values are expressed as percentages relative to the first test session (3 PM; 100%, dashed gray line). No significant differences (paired t-tests; p > 0.05) were observed between control condition (CN) and visual deprivation condition (VD). Alertness and mood showed a statistical trend toward lower levels in VD relative to CN (♦).

As shown in Table 2, sleep latency, sleep efficiency, total sleep time and N2 and N3 proportions did not differ between CN and VD experiments (also see Table 1 for an actigraphy-based comparison). VD was associated with a relative reduction in the amount of N1 sleep, a reduced REM latency, and an increased REM duration and proportion (p < 0.05, *uncorrected*). The reduced REM sleep latency in the VD condition may have been induced by a relative advance in the peak of melatonin secretion as a consequence of blindfolding, as the peak of melatonin secretion is modulated by exposure to light (Brainard et al., 2001).

**Table 2.** Sleep structure. Sleep structure in the two experimental conditions (control condition, CN; visual deprivation, VD), with group average (AVG) and standard deviation (SD). The last three columns respectively indicate the p-values, the t-scores and the effect sizes (Hedges’ g) of all contrasts across experiment. The last two rows show the relative proportion of N3 (vs. N2) sleep during the first two 20 min epochs of NREM sleep (EP1, EP2). Parameters showing a significant difference (p < 0.05, uncorrected) are marked with *. Importantly, no differences were observed in NREM-sleep parameters and in overall sleep duration and quality.

| Properties | CN Experiment |  | VD Experiment |  | Contrast |  | Effect Size |
| --- | --- | --- | --- | --- | --- | --- | --- |
|  | AVG | SD | AVG | SD | p-value | t-score | Hedges' g |
| Total Sleep Time (min) | 407.1 | 37.8 | 418.0 | 19.1 | 0.283 | 1.129 | 0.35 |
| Sleep Latency (min) | 10.0 | 8.4 | 4.7 | 2.4 | 0.057 | 2.126 | 0.83 |
| Sleep Efficiency (%) | 91.6 | 7.4 | 94.6 | 2.5 | 0.107 | 1.764 | 0.53 |
| REM Latency (min) * | 111.0 | 67.2 | 73.0 | 38.5 | 0.024 | 2.625 | 0.67 |
| Arousal Index (ar/h) | 11.4 | 7.1 | 12.7 | 7.9 | 0.317 | 1.046 | 0.17 |
| Wake After Sleep Onset (min) | 37.2 | 32.5 | 23.8 | 11.0 | 0.106 | 1.762 | 0.53 |
| N1 time (min) * | 18.4 | 16.7 | 10.7 | 6.2 | 0.043 | 2.291 | 0.59 |
| N1 proportion (%) | 4.9 | 5.0 | 2.6 | 1.6 | 0.058 | 2.137 | 0.59 |
| N2 time (min) | 222.1 | 38.5 | 215.4 | 21.6 | 0.456 | 0.773 | 0.21 |
| N2 proportion (%) | 54.6 | 8.3 | 51.7 | 6.2 | 0.171 | 1.457 | 0.38 |
| N3 time (min) | 74.9 | 28.3 | 79.6 | 25.7 | 0.585 | 0.562 | 0.17 |
| N3 proportion (%) | 18.2 | 6.3 | 18.9 | 5.7 | 0.706 | 0.392 | 0.11 |
| REM time (min) * | 91.7 | 26.2 | 112.3 | 21.9 | 0.034 | 2.425 | 0.83 |
| REM proportion (%) * | 22.3 | 5.5 | 26.8 | 4.5 | 0.042 | 2.298 | 0.86 |
| N3 in EP1 - 1<sup>st</sup> 20 min (%) | 33.8 | 23.4 | 32.1 | 21.9 | 0.826 | 0.225 | 0.07 |
| N3 in EP2 - 2<sup>nd</sup> 20 min (%) | 69.0 | 31.8 | 77.9 | 25.8 | 0.431 | 0.817 | 0.30 |

### Visual deprivation leads to lower theta activity during wakefulness in the occipital area

Previous work showed that changes in wake-dependent experience can lead to regional variations in low-frequency (*theta*) activity (Hung et al., 2013), similar to those observed in SWA during actual sleep. In line with previous findings, we found that occipital *theta* power was lower after VD than after CN (11 PM; p = 0.026, |t_(11)_| = 2.561, BCa 95% CI [0.258, 2.740]; Figure 3). This difference was no longer significant after a night of sleep (8 AM; p =0.239, |t_(11)_| = 1.246), implying a sleep-dependent re-normalization of local, experience-dependent changes in theta activity. On the other hand, no differences between CN and VD were observed in the frontal ROI, either before (p = 0.600, t_(11)_ = 0.540) or after sleep (p = 0.099, t_(11)_ = 1.802). The relative change in occipital theta activity was not correlated with the degree of reliance on visual imagery during blindfolding (r = 0.18, p = 0.572).

**Figure 3.**
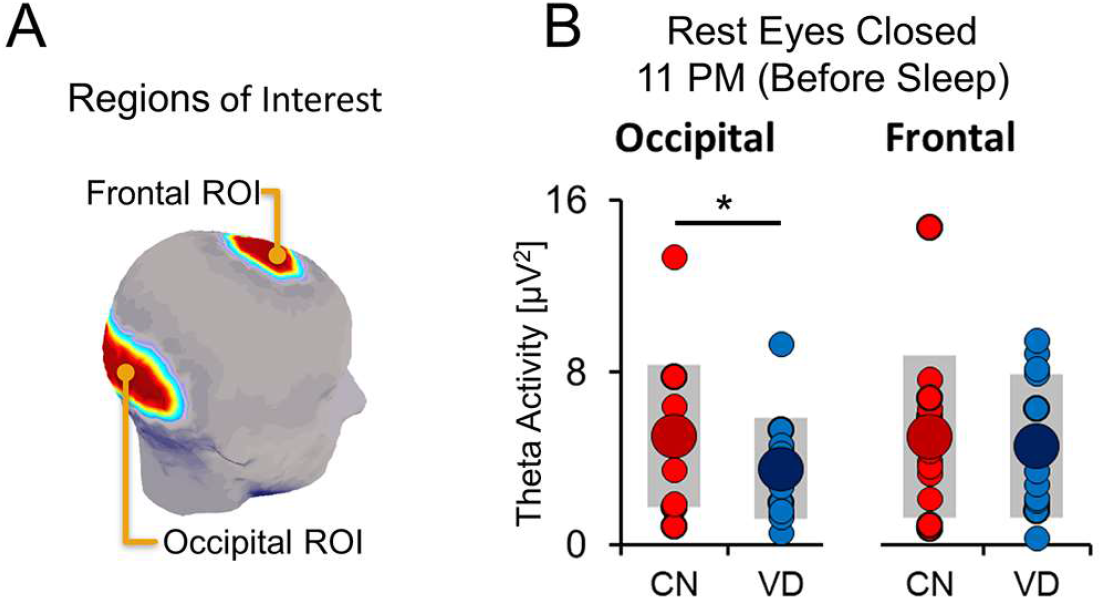
Experience-dependent changes in wake theta activity. Panel A shows the areas of the scalp covered by the occipital and the frontal regions of interest (ROI). Panel B shows variations in theta power (5-9 Hz) in control condition (CN) and after visual deprivation (VD) within the two examined ROIs (occipital, frontal). Large circles represent the group-level average, while each small circle represents a different subject (gray bars indicate one standard deviation from the mean). *, significant differences for planned comparisons (p < 0.05, paired t-test).

### Visual deprivation results in a reduction of the number of small, local occipital slow waves

During the first 20 min of NREM-sleep, SWA (Occipital: p = 0.854, |t_(11)_| = 0.189; Frontal: p = 0.240, |t_(11)_| = 1.243) and slow wave amplitude (Occipital: p = 0.518, |t_(11)_| = 0.668; Frontal: p = 0.209, |t_(11)_| = 1.334) showed no significant differences between CN and VD in the two examined ROIs (Figure 4). We also did not find any significant correlations between local changes in theta activity during wakefulness and local variations in SWA (Occipital: r = 0.11, p = 0.746; Frontal: r = −0.35, p = 0.265). There was a non-significant trend towards a lower density of slow waves in the occipital (p = 0.088, |t_(11)_| = 1.871, BCa 95% CI [−0.148, 2.604]), but not in the frontal (p = 0.606, |t_(11)_| = 0.531) ROI in the VD condition.

**Figure 4.**
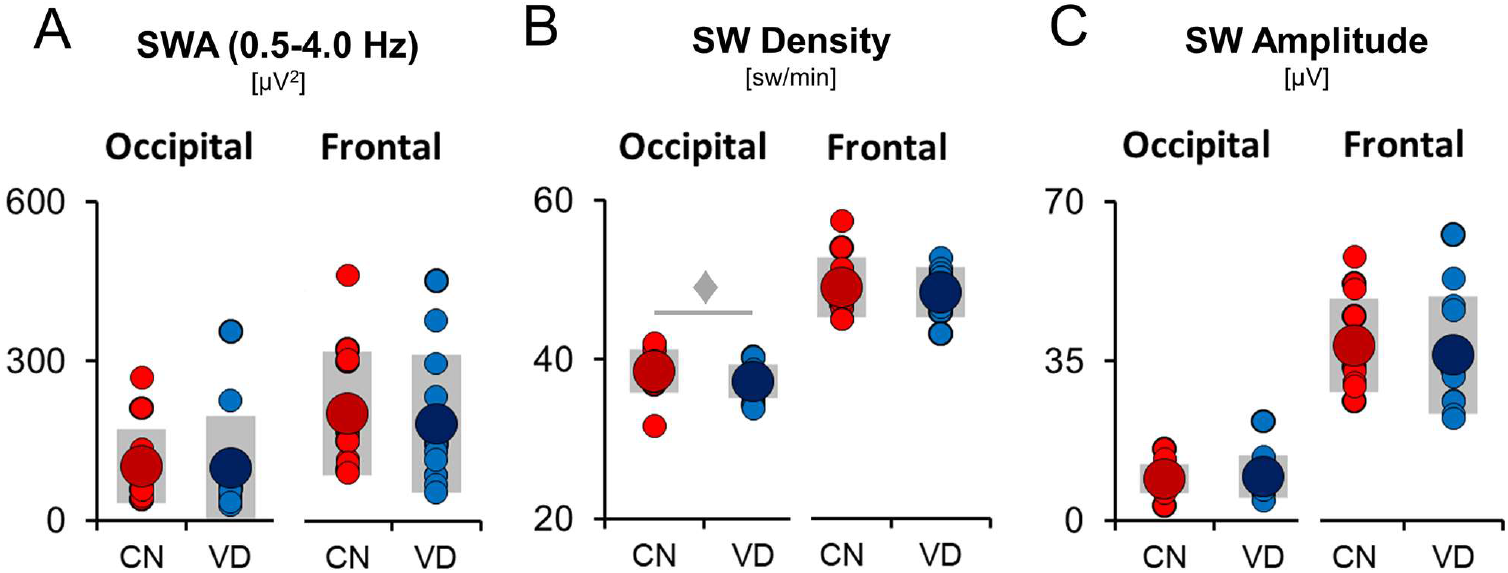
Properties of occipital and frontal slow waves. The panels show mean slow wave activity (A), slow wave density (B) and slow wave amplitude (C) in the occipital and in the frontal ROI during the first 20 min of NREM-sleep in the two experimental conditions (CN, control condition; VD, visual deprivation condition). Large circles represent the group-level average, while each small circle represents a different subject (gray bars indicate one standard deviation from the mean). ♦, nonsignificant trend (p < 0.1, paired t-test).

By examining small and large slow waves separately, we found that visual deprivation was associated with a significant decrease of small occipital slow waves (Figure 5; mean percent variation ± standard error was −8.52% ± 3.50%; p = 0.024, |t_(11)_| = 2.617, BCa 95% CI [0.940, 5.372]; Hedges’ g = 0.88), while no changes were observed for large slow waves (p = 0.197, |t_(11)_| = 1.372). Similar results were obtained using different thresholds (e.g., 1.5 or 3 MAD from the mean), or using a classification of slow waves in local (p = 0.035, |t_(11)_| = 2.405, BCa 95% CI [0.345, 3.001]; Hedges’ g = 0.64) and widespread (p = 0.449, |t_(11)_| = 0.784), based on the proportion of regional involved electrodes (*data not shown*).

**Figure 5.**
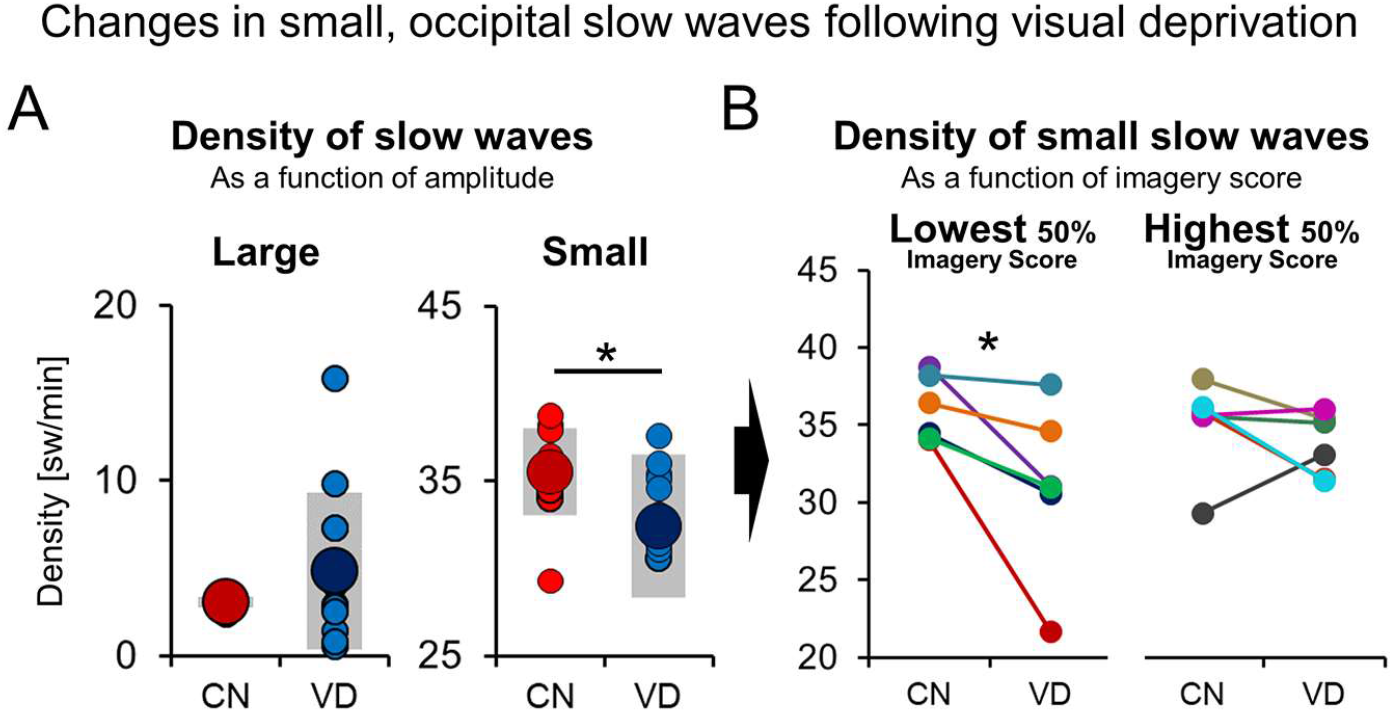
Experience-dependent changes in the density of small occipital slow waves. Visual deprivation (VD) was associated with a reduced incidence of small-amplitude waves (relative to control condition, CN), while no differences were observed for large slow waves (Panel A; *, p < 0.05, paired t-test). Large circles represent the group-level average, while each small circle represents a different subject (gray bars indicate one standard deviation from the mean). Panel B further illustrates that clear changes in the density of small waves were present in all the 6 subjects (50%) that reported the lowest reliance on visual imagery during VD, while more variability was observed in the 6 subjects who reported the highest scores. Each line/color indicates a different participant (*, p < 0.05, Wilcoxon Signed Rank Test).

### Changes in occipital sleep slow waves are modulated by visual imagery

Next, we evaluated how changes in occipital slow waves were modulated by the degree of visual imagery reported during blindfolding. We found a significant decrease in the density of small, occipital slow waves in the six (50%) participants who reported the lowest score of imagery (Wilcoxon signed rank test; p = 0.028, |z| = 2.201), but not in those who reported the highest imagery scores (p = 0.249, |z| = 1.153). A correlation analysis between the degree of visual imagery score and the overall mean amplitude showed that a low degree of visual imagery during blindfolding (VD) was associated with a relative increase in slow wave amplitude as compared to the CN condition (Figure 6A; p = 0.018, r = −0.668, BCa 95% CI [−0.208, −0.921]), likely reflecting the decreased incidence of small occipital slow waves described in Figure 5 (with no changes in large slow waves). A topographical, channel-by-channel analysis (Figure 6B) confirmed that this correlation was localized to posterior brain regions (occipital, temporal and parietal electrodes, mostly on the left side, p < 0.05, *FDR corrected*; | r | > 0.65).

**Figure 6.**
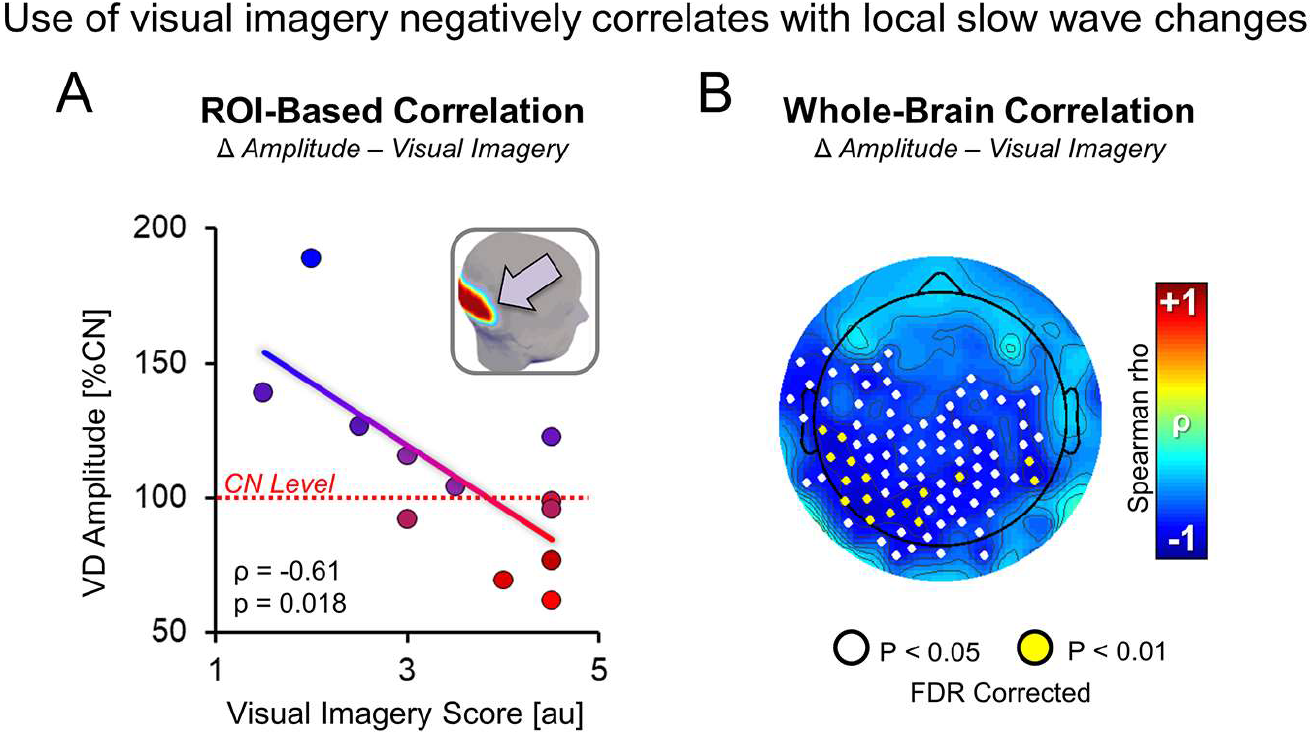
Correlation between the relative variation in slow wave amplitude after visual deprivation and reported reliance on visual imagery. Panel A shows the correlation between these parameters in the occipital ROI during the first 20 min (EP1) of NREM-sleep (p = 0.018, r = −0.61). A similar, but no longer significant correlation was observed in the second 20 min period (EP2; p = 0.189, r = −0.407). Panel B shows the results of the same correlation analysis performed on a channel-by-channel basis in EP1 (white dots indicate p < 0.05 while yellow dots indicate p < 0.01, FDR corrected).

### Imagery-related changes in slow waves involve the bilateral visual cortex

The same correlation analysis shown in Figure 6B was repeated at source level (i.e., after application of source modeling for calculation of the instantaneous current density at cortical level; Figure 7A) and confirmed a significant negative correlation (p_one-tail_ < 0.05, *FDR corrected*; | r | > 0.78; analysis restricted to negative correlations) between the visual imagery score and (0.5-4.0 Hz) signal variability (used as a proxy for slow wave amplitude in source space; see Methods) in bilateral cuneus (CUN), left posterior middle temporal gyrus (pMTG), left superior parietal lobule (SPL), left superior temporal gyrus (STG) and left inferior frontal gyrus (IFG). Based on these findings, we explored whether visual imagery also modulated the degree of inter-regional functional correlation (likely reflecting co-occurrence of slow waves) between the occipital cortex and other areas that showed a significant imagery-related modulation of slow wave amplitude. Specifically, for both CN and VD, we computed the Spearman’s correlation coefficient between the average signal of occipital voxels (CUN) and the mean time-series of pMTG, SPL, STG and IFG. Then, the relationship between the degree of visual imagery during VD and the relative strength (VD/CN ratio) of inter-regional functional couplings was investigated using the Spearman’s coefficient (two-tailed test). Results showed that reliance on visual imagery was associated with a significant increase in inter-regional functional correlation (as determined using the Spearman’s coefficient) between CUN and (Figure 7B-C) the STG (p = 0.046, r = 0.585,) and IFG (p = 0.006, r = 0.737).

**Figure 7.**
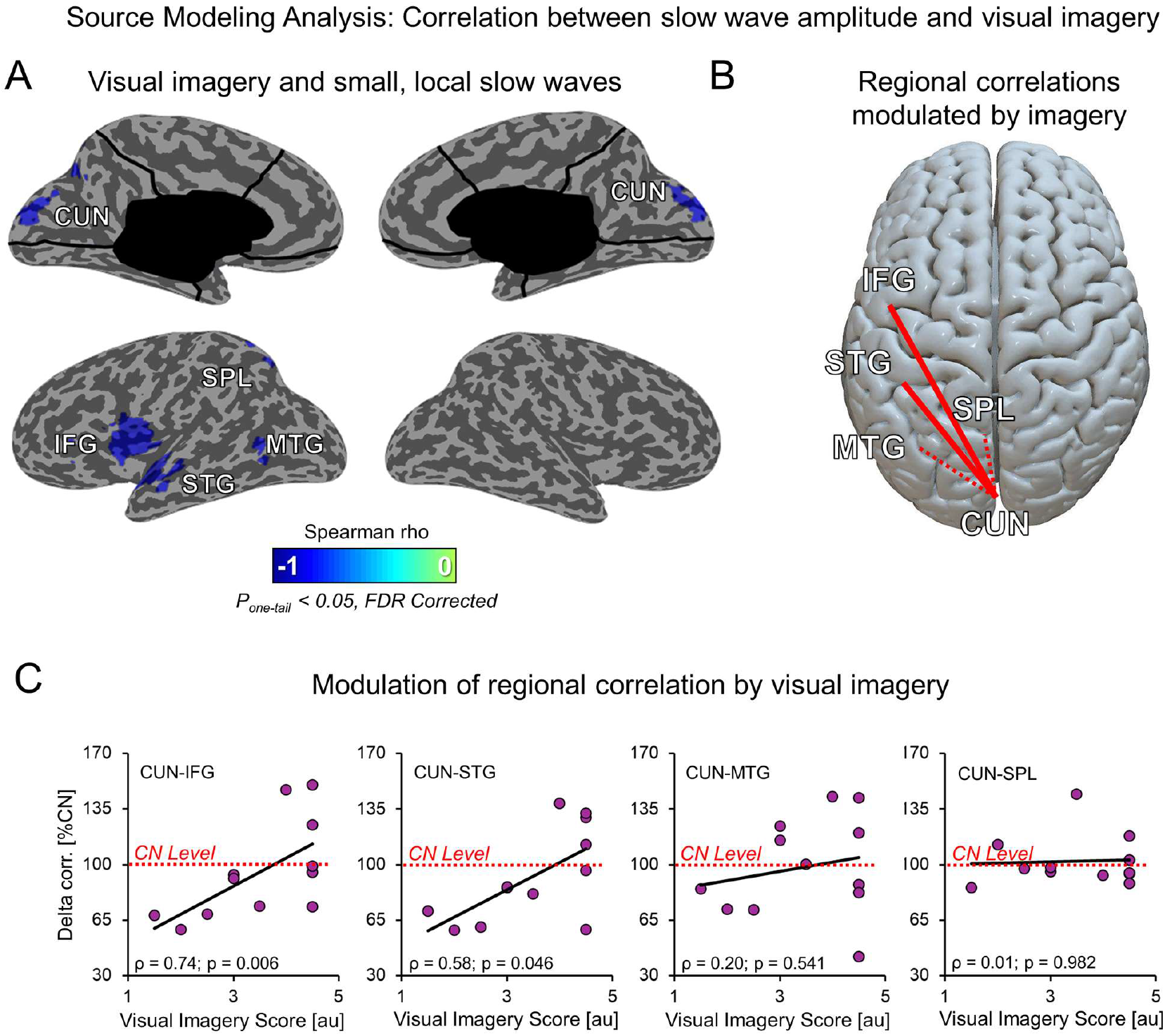
Source modeling analysis: correlation between slow wave amplitude and imagery. Panel A shows the results of the correlation between the relative variation in signal amplitude (SD of source-modeled 0.5-4.0 Hz signal) after acute visual deprivation (VD, with audiobook listening) and the reported degree of visual imagery (average of scores reported for auditory and tactile experiences) during the same time period (Spearman’s rho, pone-tail < 0.05, FDR corrected). As shown in Figure 5, the variation in amplitude reflects a change in the incidence of small amplitude, local slow waves. Panel B and C show that visual imagery also modulated the level of inter-regional correlation (Spearman’s correlation coefficient) between bilateral occipital areas (CUN) and other brain cortical regions. In panel B, red lines indicate correlations between the visual cortex and other regions that are significantly modulated by imagery (p < 0.05), while dotted lines are used for non-significant correlations. Panel C shows the scatter plots corresponding to the four examined correlations. Here the ‘CN Level’ line corresponds to the same level of inter-regional correlation observed in the control (CN) condition (100%). CUN: cuneus; STG: superior temporal gyrus; IFG: inferior frontal gyrus; MTG: middle temporal gyrus; SPL: superior temporal lobule.

To evaluate the potential association between the type of sensory experience concomitant to imagery and changes in slow wave characteristics, the same correlation analysis described above was also computed using the distinct (sub)scores of imagery provided for auditory and tactile experiences (indicating how much participants had relied on visual imagery during auditory and tactile experiences, respectively) (Figure 8). We found that the degree of reported visual imagery during auditory experiences was associated with slow wave changes similar to those observed for the analysis reported in Figure 7A. On the other hand, the degree of reliance on imagery during tactile experiences was correlated with significant slow wave activity changes in a partially overlapping network, including bilateral cuneus (CUN) and precuneus (PCUN), bilateral mid-cingulate cortex (mCC), right sensorimotor cortex and left dorsolateral prefrontal cortex (DLPFC). Importantly, in spite of the partially different sets of affected regions, correlation analyses based on auditory and tactile scores both involved a common area, that is, the bilateral occipital cortex.

**Figure 8.**
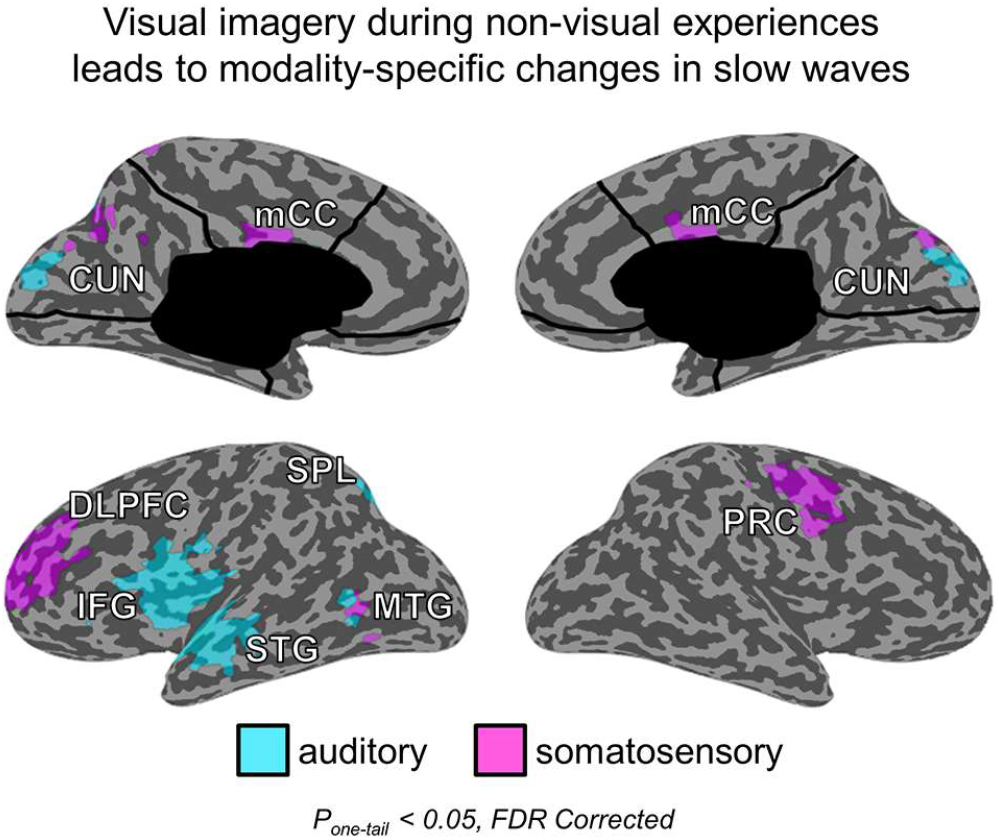
Source modeling analysis: correlation between slow wave amplitude and imagery triggered by auditory or tactile experiences. Results of the correlation between the relative variation in signal amplitude (SD of source-modeled 0.5-4.0 Hz signal) after visual deprivation (VD) and the reported exertion of visual imagery during auditory (cyan) or tactile (purple) experiences. Results indicate that participants who more strongly relied on visual imagery during tactile exploration displayed changes in slow wave activity in a network including right pre/post-central cortex, left dorsolateral prefrontal cortex and mid cingulate cortex. CUN: cuneus; STG: superior temporal gyrus; IFG: inferior frontal gyrus; MTG: middle temporal gyrus; SPL: superior temporal lobule; PRC: right pre/post-central cortex; DLPFC: dorsolateral prefrontal cortex; mCC: mid cingulate cortex.

### Visual imagery during blindfolding reduces the probability of visual hallucinations

In line with previous observations in blindfolded individuals (Merabet et al., 2004), six participants (50%) reported clear visual hallucinations (spontaneous visual perceptions projected in the external ‘objective space’ but unrelated to actual external stimuli) during the VD experiment. In most cases, hallucinations were simple (e.g., flashes of light, colored spots), short (one-to-few seconds) and appeared after at least 5-6 hours of blindfolding. However, three participants also reported brief complex hallucinations (faces, landscapes, people), and in one subject, hallucinations appeared during the first hours of visual deprivation. Interestingly, hallucinations occurred mainly in participants who reported a low reliance on visual imagery during blindfolding (Figure 9; Mann-Whitney U test, p = 0.066, z = 1.913; median score 2.75 in subjects who reported hallucinations, and 4.5 in subjects who did not report hallucinations).

**Figure 9.**
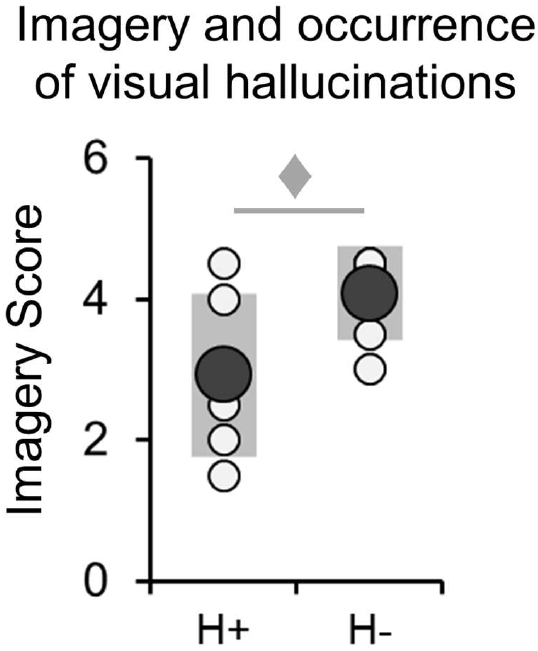
Imagery and occurrence of visual hallucinations. Six participants (50%) reported visual hallucinations during blindfolding (H+ stands for ‘hallucinations’, while H− stands for ‘no hallucinations’). Hallucinations tended to occur more often in participants who reported lower imagery scores (♦, p < 0.1, Mann-Whitney U Test).

## Discussion

By combining standardized experimental protocols and hd-EEG recordings in sleep, we demonstrated that visual deprivation during daytime induces significant changes in the number of small, local slow waves in the visual cortex during sleep. In addition, we showed that experience-dependent changes in occipital slow waves are strongly modulated by the degree of visual imagery during blindfolding, and that internally generated visual imagery may lead to network-level changes in slow wave synchronicity.

Previous work in both animal models and humans has shown that the electrophysiological marker of sleep homeostasis, NREM slow waves, can be locally modulated in an experience-dependent manner within circumscribed regions of the cerebral cortex (Huber et al., 2006, 2004; Kattler et al., 1994; Vyazovskiy et al., 2000; Wilhelm et al., 2014). However, these studies mainly targeted the sensorimotor domain, and previous attempts at investigating the relationship between experience, plasticity and sleep slow waves within the visual domain have provided inconsistent results. For instance, Miyamoto and colleagues showed that dark-rearing of cats and mice is associated with a region-specific decrease in occipital SWA during slow wave sleep (Miyamoto et al., 2003). However, the same authors noted that prolonged light deprivation (3-4 months) in adult animals had no significant effects on regional slow waves. On the other hand, in adult pigeons, short-term monocular deprivation in combination with extended wakefulness led to a homeostatic increase in SWA and in the slope of local slow waves only in the visual areas connected to the stimulated eye (Lesku et al., 2011). Finally, in humans, recent investigations based on either visual perceptual learning tasks (Bang et al., 2014; Mascetti et al., 2013) or a visual deprivation paradigm (Korf et al., 2017) failed to detect local, experience-dependent changes of slow waves in visual cortical areas. One study noted a diffuse decrease of SWA after blindfolding (Korf et al., 2017) that was prominent over fronto-parietal areas. Another study (Mascetti et al., 2013) did not find local changes in SWA but reported a positive correlation between the number of slow waves initiated in a task-related occipito-parietal area and post-sleep performance improvement.

By analyzing small-amplitude, local slow waves in the occipital cortex separately from large amplitude slow waves, we showed that 8 h of blindfolding in human volunteers who listened to audiobooks (vs. movie watching, as control condition) led to a reduction in small, local slow waves in the occipital cortex during subsequent NREM-sleep. Moreover, visual deprivation also led to a decrease in occipital *theta* activity during wakefulness – another index previously suggested to reflect local homeostatic variations in sleep need (Bernardi et al., 2015; Hung et al., 2013; Nir et al., 2017). Importantly, standard analyses of SWA or of slow wave parameters (density, amplitude) in the present study did not show any significant changes. Experience-dependent changes in sleep slow waves only emerged when small-amplitude, local slow waves were specifically examined. Several reasons could account for the finding that only smaller slow waves changed significantly in the visual system, as opposed to the sensorimotor system. In contrast to slow waves originating in centro-frontal areas, most occipital slow waves have a very small amplitude (< 10 μV) and are thus barely detectable even when no minimum amplitude threshold is used (Figure 10). Given their small amplitude, changes in slow waves originating within the visual cortex are made even more difficult to detect due to volume conduction or travelling of larger slow waves originating in other areas (Massimini et al., 2004). Indeed, the use of an amplitude threshold may favor the selection of these large, widespread slow waves at the expenses of smaller, local slow waves. Finally, we recently showed that widespread slow waves (*type I*) and local slow waves (*type II*) likely employ distinct synchronization mechanisms (subcortical for *type I* and cortical for *type II*) and are regulated differentially, with only type II waves showing homeostatic modulation (Bernardi et al., 2018; Siclari et al., 2014; Spiess et al., 2018). Hence, homeostatic changes triggered by experience-dependent plasticity should be evaluated for small, local waves independently from larger waves.

**Figure 10.**
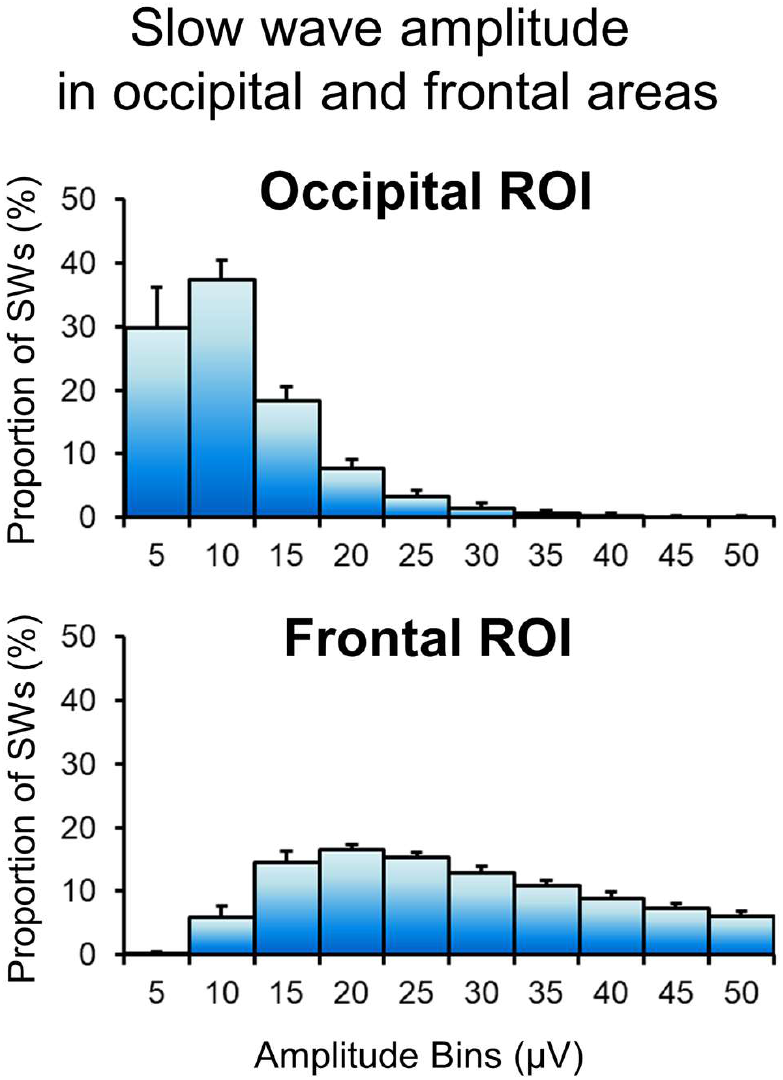
Regional differences in slow wave amplitude. Proportion of slow waves detected in the occipital ROI (left) and in the frontal (right) ROI, for different amplitude bins (5 μV steps) in the range 0-50 μV (control condition, CN). Bars represent group average, while error-bars correspond to standard error of the mean. Most occipital slow waves (~70 %) have an amplitude < 10 μV, while the minimum amplitude of most centro-frontal slow waves is typically > 5 μV. Given these premises, even a small decrease in amplitude may render local occipital waves undetectable using scalp EEG.

Importantly, slow wave changes described herein were ‘local’ in nature, and thus unlikely to reflect non-specific, global changes in sleep pressure. Indeed, visual deprivation did not induce any differences in the amount of deep sleep (N3), neither across the whole night nor the first 20 (or 40) minutes of NREM-sleep. In line with this observation, we also did not find any increases in subjective/objective sleepiness or in EEG-hallmarks known to reflect sleep need (frontal SWA in sleep and *theta* activity in wake).

The term ‘visual imagery’ is commonly used to designate internally-generated (‘*top-down*’) quasi-perceptual visual experiences occurring in the absence of a direct external stimulus (Pearson et al., 2015). A solid body of evidence indicates that visual perception and visual imagery similarly activate early and high-level visual areas (Ishai et al., 2000; Kosslyn et al., 1999). Accordingly, the content of visual imagery can be successfully decoded using a classifier that is previously trained during visual perception of the same contents (Pearson et al., 2015). We recently showed that content-specific activations in visual areas are similar between waking perception and dreaming (Siclari et al., 2017), which is perhaps the most vivid form of mental imagery that one can experience. Based on these findings, we hypothesized that visual imagery during blindfolding could mimic the effects of visual perception, thus counteracting the functional consequences of sensory deprivation. Our results are consistent with this hypothesis. Specifically, regional changes in slow wave characteristics induced by visual deprivation were strongly modulated by the degree of imagery reported by study participants: only subjects with a low reliance on visual imagery showed a clear and significant reduction of occipital sleep slow waves, while those who reported a high degree of imagery did not. Differences in the degree of visual imagery during the blindfolding period may contribute significantly to inter-subject variability and perhaps explain inconsistencies across prior studies exploring the effects of short-term visual deprivation on behavioral performance in non-visual tasks (e.g., Facchini & Aglioti, 2003; Wong et al., 2011), or on brain response to non-visual stimuli in visual areas. For instance, Lazzouni and colleagues (Lazzouni et al., 2012) reported that 6 h of visual deprivation led to clear changes in the (*cross-modal*) responsiveness of occipital areas to auditory stimuli in only half of participants. Along the same lines, these results may also have implications for our understanding of behavioral and functional heterogeneities in late blind individuals (Cattaneo et al., 2008; Cecchetti et al., 2016).

Exertion of mental imagery during non-visual experiences also led to changes in slow wave features in cortical areas related to the primarily stimulated sensory modality (e.g., language-related areas for auditory stimuli, and sensorimotor-related areas for tactile stimuli) and modulated the correlation within the 0.5-4.0 Hz frequency-band between these regions and the visual cortex during sleep. The clearest effects were observed in a left-lateralized network largely overlapping with the classical language network (Friederici & Gierhan, 2013), including middle temporal gyrus, superior temporal gyrus and inferior frontal gyrus. In this regard, it should be noted that subjects listened to audiobooks for most of the blindfolding period, and some of them spontaneously reported having used imagery to ‘visualize’ contents described in the audiobooks (other visual experiences were reported in relation to tactile perceptions, such as walking or eating during breaks). Therefore, our findings suggest that ‘internally generated’ visual experiences can not only counteract the decrease in occipital slow waves induced by lack of external inputs, but also produce changes in slow wave activity at the network level, extending beyond the visual domain.

We did not observe a correlation between imagery score and relative theta activity changes during wakefulness. Similarly, we did not observe a clear correlation between waking theta activity and sleep SWA. In fact, while both sleep slow waves and waking theta activity are thought to reflect (global) homeostatic variations in sleep need (Finelli et al., 2000; Vyazovskiy & Tobler, 2005), it is currently unclear whether these parameters also show a similar experience-dependent modulation at the local level (e.g., Hung et al., 2013). Indeed, processes such as attention or motivation are known to potentially modulate regional brain activity, and may influence the expression of low-frequency activity during wakefulness, thus representing additional sources of intra and inter-subject variability.

Half of our subjects reported the occurrence of short visual hallucinations during blindfolding that were most often elementary, but at times also complex. Similar phenomena have been reported in previous studies during blindfolding (Pitskel et al., 2007), and are known to occur in visually impaired subjects (*Charles-Bonnet syndrome;* Teunisse et al., 1996). Prolonged blindfolding may lead to an increased excitability of visual cortical areas (Boroojerdi et al., 2000; Boroojerdi et al., 2001) and may facilitate the occurrence of spontaneous visual hallucinations (Sireteanu et al., 2008). Indeed, visual hallucinations in a variety of conditions are associated with a spontaneous activation of visual cortical areas (Dickstein et al., 2004). Interestingly, in our study visual hallucinations occurred primarily in volunteers who reported a low degree of reliance on visual imagery. It is conceivable that sustained and/or reiterated exertion of visual imagery during blindfolding may partially counteract the effects of sensory deprivation on visual cortex excitability, likely through the maintenance of comparable-to-normal levels of activity in relation to internally-generated sensory experiences. In this respect, it should be noted that reported hallucinations, unlike visual imagery, were always very brief (never longer than a few seconds) and sporadic, and thus unlikely to significantly modulate plasticity and/or the excitability of visual brain areas.

## Study Limitations

While this study was performed on a relatively limited number of participants, our sample size is in line with previous work exploring the relationship between experience-dependent plasticity and slow waves in the sensorimotor domain domain (Huber et al., 2006, 2004; Kattler et al., 1994; N range =8-14), as well as with a recent study linking visual perceptual learning to slow waves originating in the occipital cortex (Mascetti et al., 2013; N=10). We believe that the strong consistency between our findings and these works, both in terms of effect size and direction, supports the validity of our observations. On the other hand, experience-dependent effects were not limited to the visual cortex, and additional studies will be needed to evaluate the spatial specificity of slow wave changes and to verify the specific role of the type of imagery on changes seen in other brain areas. Moreover, some of the reported analyses, such as the one exploring the interaction between visual imagery and hallucinations, only relied on a split-sample approach that inevitably undermined statistical power. Results of these analyses should therefore be considered as preliminary and will require verification by future studies with a larger number of participants.

Another limitation of this study is related to the use of arbitrary criteria to distinguish between local and more widespread slow waves involving the occipital region. While our results showed a good robustness under different classification methods (i.e., based on amplitude or spatial distribution) and thresholds, and are consistent with previous findings regarding the visual system (Mascetti et al., 2013), they also point to the necessity of defining more precise and reproducible approaches to distinguish slow waves based on their relative level of cortical synchronization (Bernardi et al., 2018; Siclari et al., 2014).

A direct comparison of sleep parameters between the first and second experimental condition (split-sample, Mann–Whitney U test) revealed an overall increase of N3 relative to N2 sleep in the second night (p<0.05; Table 3), consistent with a ‘first night’ effect (Agnew et al., 1966). However, we did not observe any significant order-dependent changes in N2 or N3 amount during the first 20 or 40 minutes of sleep, nor in any of the slow wave parameters in the occipital region (Table 4), suggesting that the regional distribution of experience-dependent slow wave changes is not substantially influenced by sleeping in a new environment for the first time.

**Table 3.**
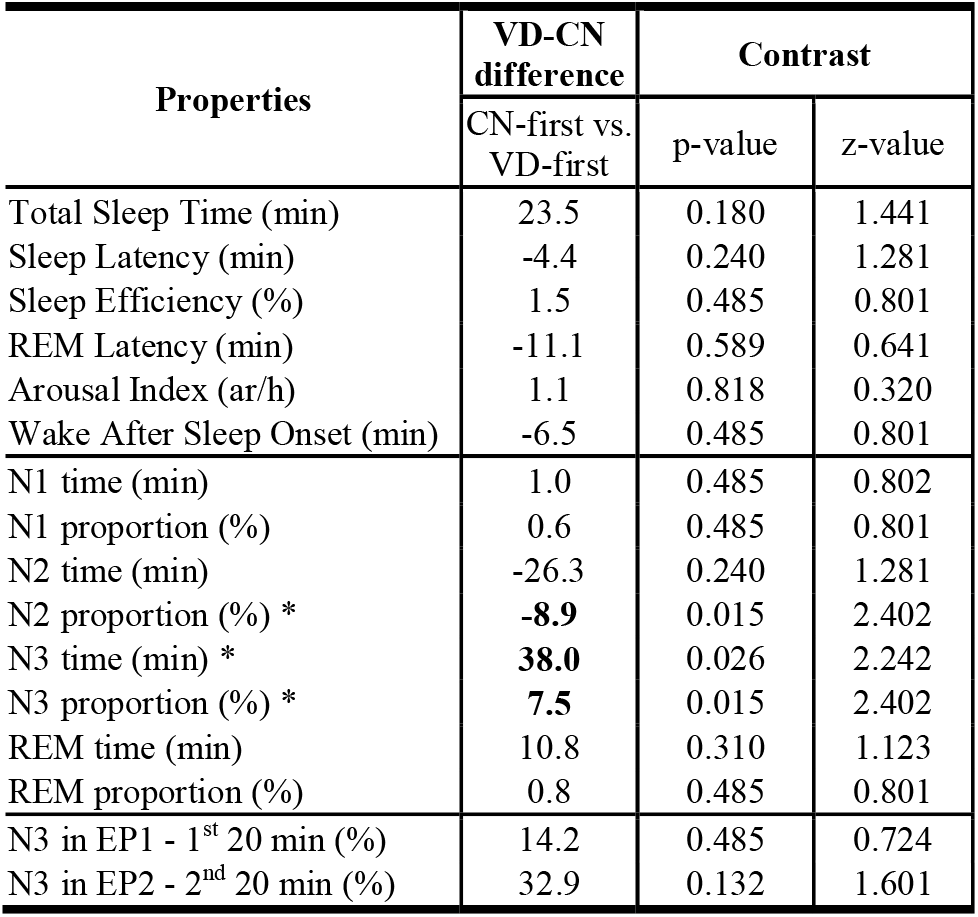
Effect of the temporal sequence of experimental conditions on VD-CN differences in sleep structure (Mann–Whitney U test for unpaired samples; N=6 in each group). Asterisks (*) mark significant effects at the p<0.05 threshold (uncorrected). The central column shows the difference in inter-condition (VD-CN) variations between subjects who performed CN first and those who performed VD first: a positive difference indicates that a higher value was observed in VD condition relative to CN condition when subjects completed the CN experiment first. VD = visual deprivation condition. CN = control night condition.

**Table 4.** Effect of the temporal sequence of experimental conditions on VD-CN differences in slow wave properties (Mann–Whitney U test for unpaired samples; N=6 in each group). No significant experiment order effects were observed. The central column shows the relative difference in inter-condition (VD-CN) variation between subjects who performed CN first and those who performed VD first: a positive difference indicates that a higher value was observed in VD condition relative to CN condition when subjects completed the CN experiment first.

| Properties (Occipital ROI) | VD-CN difference | Contrast |  |
| --- | --- | --- | --- |
|  | CN-first vs. VD-first | p-value | z-value |
| Slow wave activity (SWA) | 23.69 | 1.000 | 0.000 |
| Slow wave density | 0.27 | 1.000 | 0.000 |
| Slow wave amplitude | -0.23 | 0.818 | 0.320 |
| Density of small slow waves | 2.14 | 0.699 | -0.481 |
| Density of large slow waves | -1.87 | 0.485 | 0.801 |

## Conclusions and Future Directions

Overall, our results are consistent with the notion that imagery-related experiences can lead to functional modifications in cerebral networks and learning similar to those induced by actual perception of stimuli or execution of specific acts. More specifically, they provide preliminary evidence for how counteracting sensorimotor deprivation with learning strategies based on mental imagery, for example in rehabilitation after stroke (Dickstein et al., 2004; Sharma et al., 2009), can affect basic neural mechanisms, such as the generation of local slow waves. Moreover, they suggest that local slow wave changes may be used as a marker for imagery-related brain plastic modifications. Novel techniques allowing to modulate slow waves experimentally and locally in a noninvasive manner may open new ways for enhancing or suppressing this type of learning based on mental imagery.

## Acknowledgments

The authors thank Gianpaolo Lecciso, Ernestine Tomè, Stéphanie Dutoit, Guylaine Perron, Nadia Tobback, and Francoise Cornette for technical assistance and help with data acquisition.

